# Targeted HPLC-MS/MS Profiling of Diacylglycerols for Krill Oil Authentication and Adulteration Detection

**DOI:** 10.64898/2026.10.21.760746

**Authors:** Daria Makeeva, Louise V Michaelson

## Abstract

A rapid HPLC-MS/MS method based on multiple reaction monitoring (MRM) was developed for comprehensive analysis of diacylglycerol (DAG) molecular species in plant and marine oils. The method enabled the determination of 75 DAG molecular species within 5 min, with a 6-fold linear range and 0.02-0.2 µg/mL limits of detection. The method was successfully applied to the detection of krill oil (KO) adulteration with fish oil (FO), soybean oil (SBO), and sunflower oil (SFO). Principal component analysis (PCA) provided clear discrimination between pure and adulterated KO samples. Potential adulteration markers were identified using combined PLS-DA and random forest analysis. For FO adulteration, DAG 16:0_18:0, 20:1_20:5, 18:0_20:1, 18:0_20:0, and 20:2_22:6 were identified. For SBO and SFO adulteration DAG 18:0_18:2, 18:2_18:2, 20:2_22:6, 18:3_18:3, and 16:0_18:2 were identified as markers. The method establishes DAG profiling for quick and sensitive oil adulteration detection.

**Highlights:**

- A rapid 5-min HPLC-MS/MS-MRM method was developed for DAG profiling.
- Seventy-five DAG molecular species were quantified in plant and marine oils.
- The method enabled discrimination of pure and adulterated krill oil samples.
- DAG markers of krill oil adulteration were identified.

## 1. Introduction

Krill oil (KO) is a valuable marine-derived oil known for its high content of omega-3 fatty acids, particularly eicosapentaenoic acid (EPA) and docosahexaenoic acid (DHA), which are present in both triacylglycerol and phospholipid classes. DHA and EPA from krill and are associated with enhanced bioavailability (Ramprasath et al., 2015). Owing to its reported health benefits, including cardiovascular, cognitive, and anti-inflammatory effects, krill oil has become an increasingly important ingredient in nutritional supplements and functional foods (Kwantes & Grundmann, 2015).

Because of the rapid growth of the global KO supplement market and the substantially higher price of krill oil compared with conventional fish oil, there is a strong economic incentive for product adulteration. KO may be diluted or substituted with fish oil while maintaining comparable EPA and DHA levels, making the fraud difficult to detect. It can also be adulterated with cheap plant oils such as soybean oil (SBO) or sunflower oil (SFO) (Lee et al., 2024a). Such adulteration can change the lipid profiles of triacylglycerols (TAG), diacylglycerols (DAG), free fatty acids (FFA), or phospholipids (PL), many of which may escape detection by conventional lipid analytical techniques (Araujo et al., 2013; Cui et al., 2021a; Zailer et al., 2018).

Various analytical approaches have been developed for the detection of KO adulteration, including nuclear magnetic resonance (NMR) (Monakhova et al., 2018), Raman and infrared (IR) spectroscopy (Ahmmed et al., 2023), fluorescence spectroscopy (Ke et al., 2025), gas chromatography coupled with flame ionization detection (GC-FID) (Araujo et al., 2013; Fuller et al., 2020), thin-layer chromatography coupled with flame ionization detection (TLC-FID) (Akanbi & Barrow, 2017), and HPLC-MS/MS (Castro-Gómez et al., 2015).

HPLC–MS/MS represents a powerful analytical approach for the detection and assessment of oil adulteration, owing to its ability to provide structural characterization of individual lipid species. Since oils derived from different sources exhibit distinct fatty acyl compositions, these differences are directly reflected in the molecular profiles of their constituent lipid classes. Therefore, HPLC–MS/MS analysis offers exceptional sensitivity for detecting compositional variations and subtle modifications in lipid profiles, ensuring the authenticity of oil-based products.

Existing HPLC–MS methods are predominantly focused on the analysis of major lipid classes, including TAG (Araujo et al., 2018) and PL (Cui et al., 2021b). This is unsurprising, as these lipid classes are generally the most abundant and are highly studied. However, the lipid composition of KO is very complex, with TAG and PL fractions exhibiting extensive molecular diversity due to the presence of numerous fatty acyl chains, including 14:0, 16:0, 16:1n7, 18:0, 18:1n9, 18:1n7, 18:2n6, 18:4n3, 20:5n3, and 22:6n3 (Akanbi & Barrow, 2017b). Consequently, the comprehensive characterization of lipid molecular species within these major lipid classes often requires lengthy chromatographic separations, high-resolution instrumentation, and optimized methods capable of resolving closely related lipid species.

In krill, as in other organisms, diacylglycerol (DAG) represents a central intermediate in glycerolipid metabolism, serving as a key precursor for the biosynthesis of both TAG and PL via the Kennedy pathway(Gibellini & Smith, 2010). TAG biosynthesis occurs through the enzymatic acylation of DAG resulting in the formation of storage lipids. Similarly, DAG contributes to PL biosynthesis, where it undergoes head-group modification to generate major membrane lipids such as phosphatidylcholine (PC) and phosphatidylethanolamine (PE). DAG connects membrane synthesis and energy storage and occupies a central position in glycerolipid metabolism.

DAG profiling for the detection of oil adulteration has received considerably less attention than TAG-(Akanbi & Barrow, 2017b) and PL-based approaches (Cui et al., 2021c). DAGs represent a promising and potentially underexploited lipid class for oil authentication. Compared with TAG, the number of possible DAG molecular species is substantially lower owing to the presence of only two fatty acid moieties rather than three. This enables a more focused and comprehensive characterization of the DAG fraction with reduced analytical complexity and significantly shorter analysis times. Importantly, DAGs occupy a central position in glycerolipid metabolism and can retain compositional characteristics associated with both membrane phospholipids and storage triacylglycerols. Consequently, DAG profiling has the potential to capture complementary information on lipid composition while providing a simpler and more cost-effective analytical strategy for oil authentication and adulteration detection.

In this study, we developed a rapid and sensitive DAG-based method for the detection of KO adulteration with FO and other plant-derived oils, including SBO and SFO. The method combines targeted MRM analysis with response factor-based quantitation, enabling accurate profiling of DAG species within a 5 min HPLC-MS/MS run. The approach proved effective for the detection and quantification of oil adulteration.

## 2. Materials and methods

### 2.1 Chemical reagents

LC-MS grade acetonitrile, isopropanol, formic acid, methanol, and HPLC-grade hexane, chloroform, and formic acid were acquired from Thermo Fisher Scientific (Waltham, MA, United States). A 10 M ammonium formate solution was obtained from Sigma-Aldrich (Buchs, Switzerland).

All DAG standards were obtained from Nu-Chek Prep (Elysian, Minnesota, USA). The following abbreviations, which indicate the length of each acyl chain and the number of double bonds per chain, were used in this study: dipalmitin (DAG 16:0_16:0), distearin (DAG 18:0_18:0), diarachidin (TAG 20:0_20:0), diolein (DAG 18:1_18:1Δ9), dilinolein (DAG 18:2_18:2Δ9,12), dilinolenin (DAG 18:3_18:3Δ9,12,15), dieicosenoin (DAG 20:1_20:1Δ11), dieicosadienoin (DAG 20:2_20:2Δ11,14), dieicosatrienoin (DAG 20:3_20:3Δ11,14,17), diarachidonin (DAG 20:4_20:4Δ5,8,11,14), didocosahexaenoin (DAG 22:6_22:6 Δ4,7,10,13,16,19).

Fish oil (FO), soybean oil (SBO), and sunflower oil (SFO) pharmaceutical secondary standards (Supelco), together with a krill oil United States Pharmacopeia (USP) reference standard, were obtained from Sigma-Aldrich (Buchs, Switzerland). Artificial adulterated samples were prepared by gravimetrically mixing the USP krill oil reference standard with pharmaceutical secondary standards of fish, soybean, and sunflower oils at predefined proportions.

### 2.2 HPLC-MS/MS analysis conditions

LC-MS/MS analyses were performed using a SCIEX ExionLC AD UHPLC system (SCIEX, USA) coupled to a SCIEX 6500+ triple quad liquid MS system. DAGs were ionized using electrospray ionization (ESI) and detected in positive ion mode using MRM. Chromatographic separations were performed on an Zorbax Eclipse Plus C18 RRHD column (50 × 2.1 mm, 1.8 µm, Agilent). Mobile phase A consisted of acetonitrile:water (50:50, v/v) with 10 mM ammonium formate and 0.1% formic acid, while mobile phase B consisted of isopropanol:acetonitrile:water (88:10:2, v/v/v) with 2 mM ammonium formate and 0.02% formic acid. The flow rate was 0.6 mL/min. The injection volume was 1 µL, the column oven was maintained at 40 °C, and the autosampler was set at 10 °C. A needle wash with isopropanol was performed before each injection, using a wash time of 2 seconds and a wash speed of 10 µL/s. The total run time was 5.0 minutes with the following solvent gradient: 0.0 min, 70% B; 0.1 min, 70% B; 3.5 min, 95% B; 3.6 min, 99% B; 4.0 min, 99% B; 4.1 min, 70% B; 5.0 min, stop run.

The ESI parameters curtain gas (CUR), gas 1 (GS1), and gas 2 (GS2) were set to 35, 50, and 50 psi, respectively. Optimal parameters for Ion Spray Voltage (IS), Collision gas (CAD), and Temperature (TEM) were 4500 V, low and 200°C, respectively. Mass spectra was acquired using scheduled MRM with a cycling time of one second and an MRM window of 60 seconds. MRM values for individual lipid species are shown in Table 1. Declustering (DP), entrance (EP), cell exit potentials (CXP), and collision energy (CE) were optimising using standards and were 100, 10, 30 and 10, respectively.

**Table 1.** DAG molecule transitions information for the MRM MS/MS quantitation as well as retention time and RFs.

| Molecule | Precursor<br>m/z | Precursor<br>Adduct | Product 1<br>m/z | Product 2<br>m/z | RT, min | RF |
| --- | --- | --- | --- | --- | --- | --- |
| DAG 16:0_18:1 | 612.6 | [M+NH<sub>4</sub>] | 339.3 | 313.3 | 2.3 | 0.65 |
| DAG 16:0_18:2 | 610.5 | [M+NH<sub>4</sub>] | 337.3 | 313.3 | 1.9 | 0.48 |
| DAG 18:0_18:1 | 640.6 | [M+NH<sub>4</sub>] | 339.3 | 341.3 | 2.8 | 0.77 |
| DAG 18:0_18:2 | 638.6 | [M+NH<sub>4</sub>] | 337.3 | 341.3 | 2.4 | 0.60 |
| DAG 18:0_18:3 | 636.6 | [M+NH<sub>4</sub>] | 335.3 | 341.3 | 2.1 | 0.49 |
| DAG 18:1_18:2 | 636.6 | [M+NH<sub>4</sub>] | 337.3 | 339.3 | 2.0 | 0.83 |
| DAG 18:1_18:3 | 634.5 | [M+NH<sub>4</sub>] | 335.3 | 339.3 | 1.7 | 0.72 |
| DAG 16:0_20:5 | 632.5 | [M+NH<sub>4</sub>] | 359.3 | 313.3 | 1.6 | 0.30 |
| DAG 18:0_20:5 | 660.6 | [M+NH<sub>4</sub>] | 359.3 | 341.3 | 2.0 | 0.42 |
| DAG 16:0_22:6 | 658.5 | [M+NH4] | 385.3 | 313.3 | 1.7 | 0.29 |
| DAG 18:2_20:5 | 656.5 | [M+NH4] | 359.3 | 337.3 | 1.3 | 0.48 |
| DAG 18:3_20:5 | 654.5 | [M+NH4] | 359.3 | 335.3 | 1.1 | 0.37 |
| DAG 20:5_20:5 | 678.5 | [M+NH4] | 359.3 | — | 1.0 | 0.31 |
| DAG 20:1_20:5 | 686.6 | [M+NH4] | 359.3 | 367.3 | 1.9 | 0.93 |
| DAG 18:1_22:6 | 684.6 | [M+NH4] | 385.3 | 339.3 | 1.8 | 0.64 |
| DAG 16:0_18:3 | 608.5 | [M+NH4] | 335.3 | 313.3 | 1.6 | 0.37 |
| DAG 18:1_20:5 | 658.5 | [M+NH4] | 359.3 | 339.3 | 1.6 | 0.65 |
| DAG 18:2_22:6 | 682.5 | [M+NH4] | 385.3 | 337.3 | 1.4 | 0.47 |
| DAG 18:3_22:6 | 680.5 | [M+NH4] | 385.3 | 335.3 | 1.2 | 0.36 |
| DAG 16:1_18:1 | 610.5 | [M+NH4] | 339.3 | 311.3 | 1.9 | 0.65 |
| DAG 16:2_18:1 | 608.5 | [M+NH4] | 339.3 | 309.2 | 1.5 | 0.65 |
| DAG 16:1_18:2 | 608.5 | [M+NH4] | 337.3 | 311.3 | 1.5 | 0.48 |
| DAG 16:1_20:5 | 630.5 | [M+NH4] | 359.3 | 311.3 | 1.2 | 0.30 |
| DAG 16:2_20:5 | 628.5 | [M+NH4] | 359.3 | 309.2 | 1.0 | 0.30 |
| DAG 20:4_20:5 | 680.5 | [M+NH4] | 359.3 | 361.3 | 1.1 | 0.31 |
| DAG 20:5_22:6 | 704.5 | [M+NH4] | 385.3 | 359.3 | 1.1 | 0.29 |
| DAG 14:0_22:6 | 630.5 | [M+NH4] | 385.3 | 285.2 | 1.3 | 0.29 |
| DAG 16:1_22:6 | 656.5 | [M+NH4] | 385.3 | 311.3 | 1.3 | 0.29 |
| DAG 16:2_22:6 | 654.5 | [M+NH4] | 385.3 | 309.2 | 1.2 | 0.29 |
| DAG 18:4_22:6 | 678.5 | [M+NH4] | 385.3 | 333.2 | 1.0 | 0.36 |
| DAG 20:0_22:6 | 714.6 | [M+NH4] | 385.3 | 369.3 | 2.3 | 0.56 |
| DAG 20:4_22:6 | 706.5 | [M+NH4] | 385.3 | 361.3 | 1.4 | 0.65 |
| DAG 14:0_16:4 | 550.4 | [M+NH4] | 305.2 | 285.2 | 0.8 | 0.30 |
| DAG 16:0_16:4 | 578.5 | [M+NH4] | 305.2 | 313.3 | 1.1 | 0.30 |
| DAG 16:4_18:1 | 604.5 | [M+NH4] | 339.3 | 305.2 | 1.1 | 0.65 |
| DAG 16:4_20:0 | 634.5 | [M+NH4] | 369.3 | 305.2 | 1.5 | 0.57 |
| DAG 16:4_20:5 | 624.5 | [M+NH4] | 359.3 | 305.2 | 0.7 | 0.30 |
| DAG 14:0_14:0 | 530.5 | [M+NH4] | 285.2 | — | 1.3 | 0.30 |
| DAG 16:0_18:4 | 606.5 | [M+NH4] | 333.2 | 313.3 | 1.4 | 0.37 |
| DAG 14:0_16:1 | 556.5 | [M+NH4] | 311.3 | 285.2 | 1.3 | 0.30 |
| DAG 14:0_18:0 | 586.5 | [M+NH4] | 341.3 | 285.2 | 2.2 | 0.42 |
| DAG 14:0_18:1 | 584.5 | [M+NH4] | 339.3 | 285.2 | 1.8 | 0.65 |
| DAG 14:0_18:2 | 582.5 | [M+NH4] | 337.3 | 285.2 | 1.5 | 0.48 |
| DAG 14:0_18:3 | 580.5 | [M+NH4] | 335.3 | 285.2 | 1.2 | 0.37 |
| DAG 14:0_18:4 | 578.5 | [M+NH4] | 333.2 | 285.2 | 1.0 | 0.37 |
| DAG 14:0_20:0 | 614.6 | [M+NH4] | 369.3 | 285.2 | 2.8 | 0.57 |
| DAG 14:0_20:5 | 604.5 | [M+NH4] | 359.3 | 285.2 | 1.1 | 0.30 |
| DAG 14:0_16:2 | 554.5 | [M+NH4] | 309.2 | 285.2 | 1.1 | 0.30 |
| DAG 16:1_18:4 | 604.5 | [M+NH4] | 333.2 | 311.3 | 1.1 | 0.37 |
| DAG 18:1_18:4 | 632.5 | [M+NH4] | 333.2 | 339.3 | 1.4 | 0.72 |
| DAG 18:4_20:0 | 662.6 | [M+NH4] | 369.3 | 333.2 | 2.0 | 0.65 |
| DAG 16:0_16:0 | 586.5 | [M+NH4] | 313.3 | — | 2.2 | 0.30 |
| DAG 18:4_20:5 | 652.5 | [M+NH4] | 359.3 | 333.2 | 0.9 | 0.37 |
| DAG 18:0_18:0 | 642.6 | [M+NH4] | 341.3 | — | 3.1 | 0.54 |
| DAG 18:1_18:1 | 638.6 | [M+NH4] | 339.3 | — | 2.3 | 1.00 |
| DAG 18:2_18:2 | 634.5 | [M+NH4] | 337.3 | — | 1.6 | 0.66 |
| DAG 18:3_18:3 | 630.5 | [M+NH4] | 335.3 | — | 1.1 | 0.44 |
| DAG 20:0_20:0 | 698.7 | [M+NH4] | 369.3 | — | 3.9 | 0.85 |
| DAG 20:1_20:1 | 694.6 | [M+NH4] | 367.3 | — | 3.2 | 1.56 |
| DAG 20:2_20:2 | 690.6 | [M+NH4] | 365.3 | — | 2.5 | 1.02 |
| DAG 20:3_20:3 | 686.6 | [M+NH4] | 363.3 | — | 1.9 | 0.56 |
| DAG 20:4_20:4 | 682.5 | [M+NH4] | 361.3 | — | 1.3 | 0.31 |
| DAG 22:6_22:6 | 730.5 | [M+NH4] | 385.3 | — | 1.2 | 0.28 |
| DAG 20:2_22:6 | 710.6 | [M+NH4] | 385.3 | 365.3 | 1.9 | 0.65 |
| DAG 14:0_16:0 | 558.5 | [M+NH4] | 313.3 | 285.2 | 1.7 | 0.30 |
| DAG 16:0_16:1 | 584.5 | [M+NH4] | 311.3 | 313.3 | 1.8 | 0.30 |
| DAG 16:0_16:2 | 582.5 | [M+NH4] | 309.2 | 313.3 | 1.5 | 0.48 |
| DAG 16:0_16:3 | 580.5 | [M+NH4] | 307.2 | 313.3 | 1.3 | 0.37 |
| DAG 16:0_18:0 | 614.6 | [M+NH4] | 341.3 | 313.3 | 2.7 | 0.42 |
| DAG 18:0_20:0 | 670.6 | [M+NH4] | 369.3 | 341.3 | 3.4 | 0.69 |
| DAG 18:1_20:0 | 668.6 | [M+NH4] | 369.3 | 339.3 | 2.9 | 0.92 |
| DAG 18:0_20:1 | 668.6 | [M+NH4] | 367.3 | 341.3 | 3.1 | 1.05 |
| DAG 18:2_20:0 | 666.6 | [M+NH4] | 369.3 | 337.3 | 2.5 | 0.76 |
| DAG 18:1_20:1 | 666.6 | [M+NH4] | 367.3 | 339.3 | 2.8 | 1.28 |
| DAG 18:3_20:0 | 664.6 | [M+NH4] | 369.3 | 335.3 | 2.2 | 0.65 |

### 2.3. Calibration curves and quantitation of DAGs by response factor

Calibration curves were built for each of the DAG standards listed above (Chemical reagents). These were dissolved in a CHCl_3_/MeOH (2:1, v/v) mixture to prepare three 10 mg/mL stock solutions for each DAG standard. The stock solutions were stored at −20 °C. Prior to use, the solutions were thawed to room temperature, and three sets of standard mixtures were prepared by combining stock solutions and diluting them with mobile phase B, yielding working solutions with the following concentrations of each lipid: 0.0001, 0.010, 0.05, 0.10, 0.50, 1.00, 2.50, 5.00 and 10.00 µg/mL. The average peak areas were used to generate the final calibration curves.

The calibration curves were used to calculate response factors (RFs) by the approach described by (Holčapek et al., 2005). Briefly, the slope of each individual calibration curve was used to estimate response factors for single fatty acids DAGs in relation to DAG 18:1_18:1, the response factor of which was set to 1:

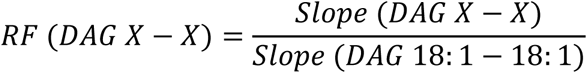

The RFs of mixed fatty acid DAGs were calculated as the average of the corresponding RFs of individual fatty acid DAGs:

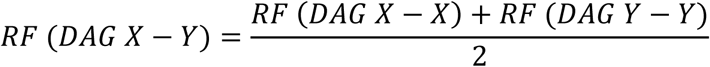

### 2.4 Oil sample preparation and artificial adulteration experiments for HPLC-MS/MS analyses

Standard oil samples for DAG profile analysis of the pure oils were subjected to a 100-fold dilution prior to analysis using CHCl_3_/MeOH (2:1, v/v), resulting in an approximate concentration of 10 mg/mL. Adulterated samples were prepared by mixing 10 mg/mL oil solutions at appropriate volume ratios to obtain final adulterant concentrations of 2.5, 4, 5, 10, and 15% (w/w). Prior to analysis, samples were diluted by mixing 10 µL of the oil sample (pure or adulterated) with 10 µL of a 2 µg/mL DAG 20:3/20:3 internal standard solution and 80 µL of mobile phase A. The resulting mixtures were subsequently subjected to DAG analysis.

### 2.5 Data processing and statistics

Data processing was performed using *Skyline* (Adams et al., 2020) software, which provided good visualization of the MS/MS results. For each target DAG molecule, product ion lists were generated using an *in silico* library created with *LipidCreator* (Peng et al., 2020a).

Comparative study of DAG composition of oils was carried out using partial least squares discriminate analysis (PLS−DA), and variable importance in projection (VIP) scores and heat map visualization, in MetaboAnalyst 6.0 (Pang et al., 2024). Log transformation (base 10) and data scaling (mean-centering and division by the square root of each variable’s standard deviation) were performed prior to statistical analysis.

To identify molecular markers associated with adulteration, PLS-DA and random forest classification models were applied. The top 10 discriminating compounds ranked according to VIP scores from PLS-DA and variable importance values from random forest were compared. Five compounds identified by both complementary approaches were selected as robust candidate markers. These markers were subsequently validated using one-way ANOVA to confirm significant differences among the sample groups.

## 3. Results and discussion

### 3.1 HPLC-MS/MS Method Development for DAG Quantification: Chromatographic and Mass Spectrometric Optimization, Response Factor Determination, and Method Validation

In positive-ion MS/MS, DAG is preferentially ionized as [M+NH₄]⁺ or [M+Na]⁺ precursor ions and typically fragment through cleavage of the ester bonds, resulting in characteristic neutral losses of fatty acids (Figure 1). These neutral losses generate monoacylglycerol (MAG)-type fragment ions that enable identification of the fatty acyl chains present in the DAG species. Intact DAG precursor ions also undergo dehydration, producing [M+NH_3_−H₂O]⁺ fragment ions, which are typically the most abundant product ions under mild collision-induced dissociation (CID) conditions. However, different DAG species with distinct fatty acyl compositions can produce identical dehydration fragments, increasing the risk of misidentification and fragment overlap. Therefore, only MAG-type fragment ions were used for DAG identification (Table 1). In this approach, the detection of two MAG-type fragment ions originating from the same precursor ion [M+NH_4_]^+^ was considered evidence for the presence of the corresponding DAG species in the sample.

**Figure 1.**
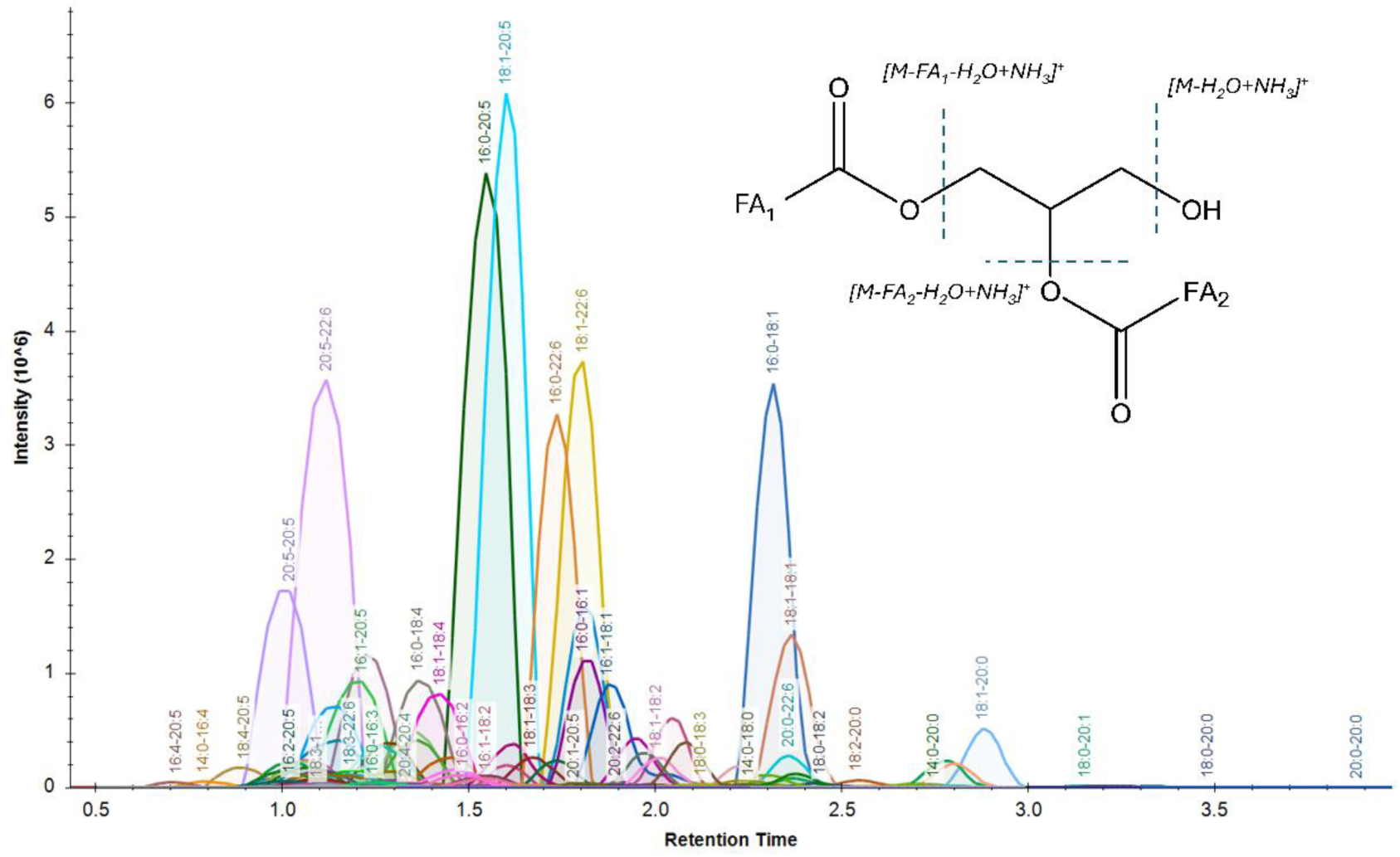
HPLC-MS/MS multiple reaction monitoring (MRM) chromatogram of diacylglycerols (DAGs) detected in KO using the transitions listed in Table 1. Separation was performed using mobile phase A (acetonitrile:water, 50:50, v/v, containing 10 mM ammonium formate and 0.1% formic acid) and mobile phase B (isopropanol:acetonitrile:water, 88:10:2, v/v/v, containing 2 mM ammonium formate and 0.02% formic acid) at a flow rate of 0.6 mL min⁻¹. The gradient program was: 0.0-0.1 min, 70% B; 3.5 min, 95% B; 3.6-4.0 min, 99% B; 4.1 min, 70% B; and 5.0 min, end of run.

The list of DAG targets was generated using an *in silico* library created with *LipidCreator* (Peng et al., 2020b). All predicted precursor and product ions, except for the dehydration [M+H−H₂O]⁺ fragments, were screened in pooled sample containing equal amounts of KO, FO, SBO, and SFO. Only targets yielding both expected MAG-type product ions were retained for subsequent analysis. The ESI parameters were optimized for ionization, with final IS, CAD, and TEM values set to 4500 V, low, and 200 °C, respectively. DP, EP, CXP, and CE were optimized using standards to achieve the highest signal intensity and were set to 100, 10, 30, and 10, respectively.

Different chromatographic parameters, including column temperature, mobile phase composition, and gradient profile, were evaluated to achieve optimal separation while maintaining the shortest possible analysis time. The final chromatographic conditions are described in Section 2.2. A column temperature of 40°C resulted in lower backpressure, allowing an increased flow rate and consequently reducing the overall analysis time.

For each identified DAG species, retention time (RT) evaluation was used to support structural assignment. Similar to TAG (Makeeva et al., 2026), the RT of DAG in reverse-phase chromatography depends on the total carbon number (CN) and the number of double bonds (DB) in the fatty acyl chains. An increase in the total carbon number results in a positive shift in RT, whereas each additional double bond contributes to a decrease in RT. Therefore, when DAG species identified based on MS/MS fragment alignment are arranged according to the same total carbon number and ordered by DB increase, a gradual decrease in retention time should be observed (Figure 2). The position of each DAG species within this retention time trend provides an additional confirmation of the assigned structure.

**Figure 2.**
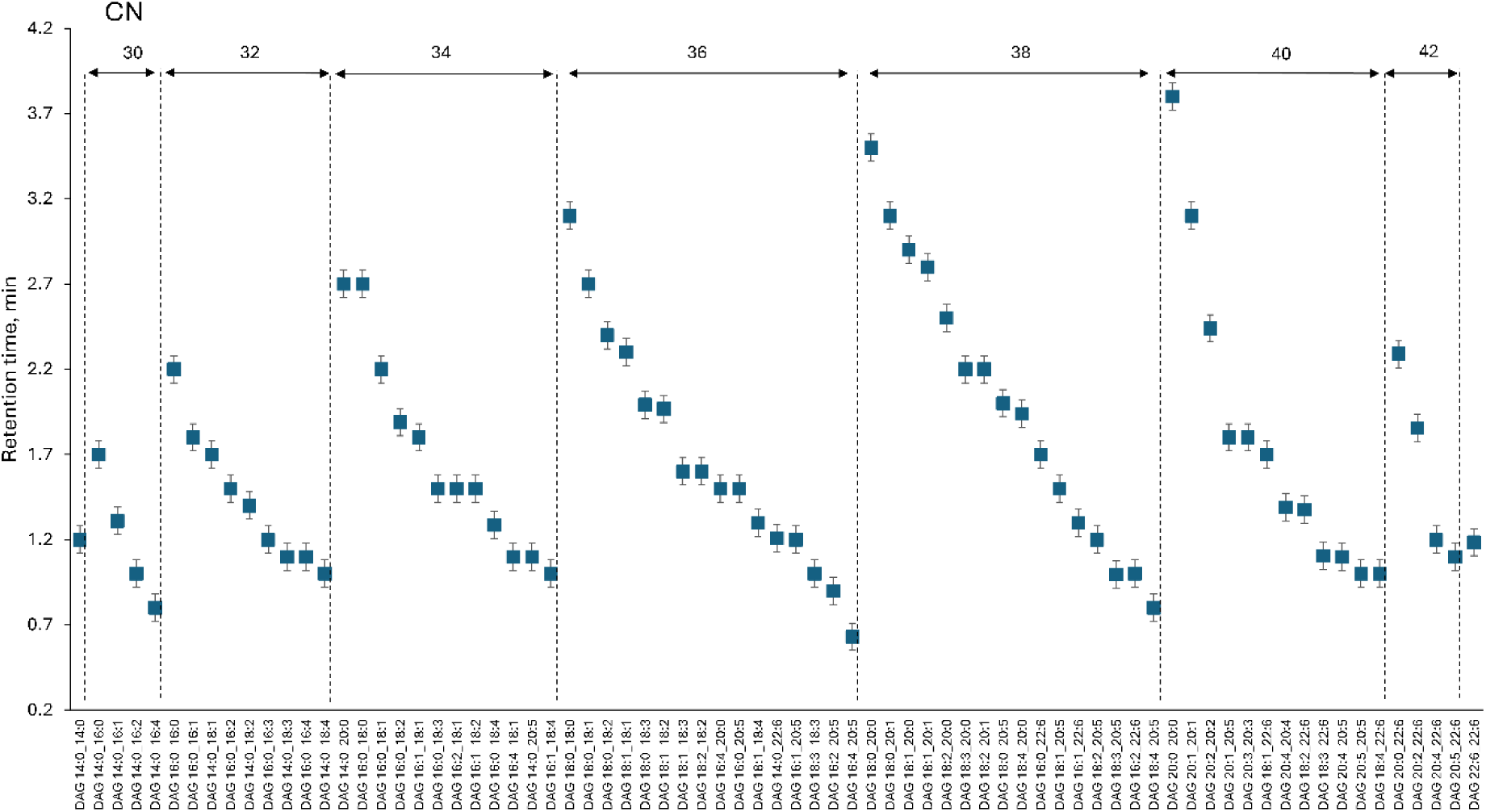
Retention times of the DAG species in a pooled sample containing KO, FO, SBO, and SFO. RT of DAG, grouped by total carbon number and increasing desaturation.

**Figure 3.**
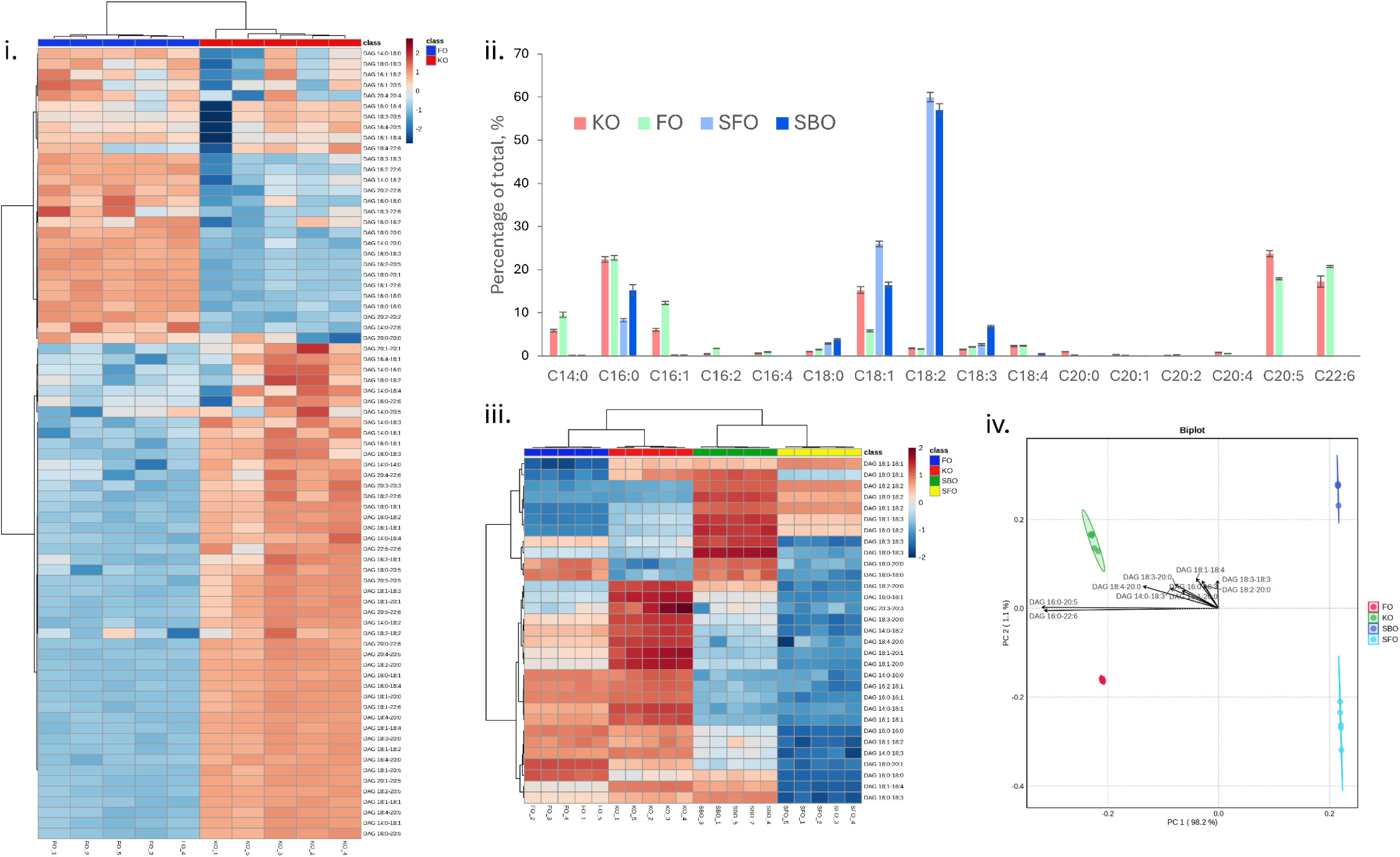
DAG composition of pure KO, FO, SFO, and SBO oils. (i) Heatmap of selected DAG molecular species identified and quantified in KO and FO. Simplified for visual clarity. (ii) Fatty acyl composition of the DAG fraction of the analysed pure oils. (iii) Heatmap of DAG molecular species detected across KO, FO, SFO, and SBO. (iv) PCA biplot of the pure oils based on DAG profiles.

Thus, a method for the HPLC-MS/MS determination of 75 DAG species within 5 minutes was developed, comprehensively covering the DAG molecular species present in KO and its most common adulterants and providing profiling of the oils (Figure 1).

The different MS responses of DAG species, which depend on their fatty acyl compositions (Holčapek et al., 2005), (Makeeva et al., 2026), require the application of response factors (RFs) for accurate quantitation, particularly when detecting adulterated samples. To account for the influence of fatty acyl composition, RFs were calculated using authentic standards of DAG species containing single fatty acids. Subsequently, individual RFs for mixed fatty acid DAG species were calculated by averaging the corresponding RFs of the single fatty acid DAG standards, as described in *Section 2.3*. For each sample analysed the RFs were used to produce the quantitative data (*Suppl. Materials 1, Table 1-4*).

The method was validated using authentic standards of DAG species containing identical fatty acyl chains. Calibration curves were constructed for these standards (Section 2.2), and the linear range, LOD, and LOQ were determined. The method exhibited a 6-fold linear range, with correlation coefficients over 0.98 and LOQ values ranging from 0.06 to 0.3 µg/mL (Table 2).

**Table 2.** Method validation and response factors (RFs).

| DAG | a | b | R<sup>2</sup> | Linear range,<br>ng/mL | RFs | LOD,<br>µg/mL | LOQ,<br>µg/mL |
| --- | --- | --- | --- | --- | --- | --- | --- |
| DAG 16:0_16:0 | 6287 | 1724461 | 0.99106 | 0.3-10000.0 | 0.295 | 0.1 | 0.3 |
| DAG 18:0_18:0 | 11511 | 5547642 | 0.98040 | 0.3-10000.0 | 0.541 | 0.1 | 0.3 |
| DAG 18:1_18:1 | 21292 | 7747368 | 0.98408 | 0.1-10000.0 | 1.000 | 0.033 | 0.1 |
| DAG 18:2_18:2 | 14114 | 1496555 | 0.99916 | 0.2-10000.0 | 0.663 | 0.066 | 0.2 |
| DAG 18:3_18:3 | 9458 | -33243 | 0.99521 | 0.3-10000.0 | 0.444 | 0.1 | 0.3 |
| DAG 20:0_20:0 | 18049 | 8028354 | 0.97575 | 0.30-10000.0 | 0.848 | 0.1 | 0.3 |
| DAG 20:1_20:1 | 33254 | 18519345 | 0.98427 | 0.06-10000.0 | 1.562 | 0.02 | 0.06 |
| DAG 20:2_20:2 | 21713 | 8985790 | 0.98174 | 0.06-10000.0 | 1.020 | 0.02 | 0.06 |
| DAG 20:3_20:3 | 12011 | 1627111 | 0.99845 | 0.1-10000.0 | 0.564 | 0.1 | 0.3 |
| DAG 20:4_20:4 | 6499 | 472317 | 0.99723 | 0.1-10000.0 | 0.305 | 0.1 | 0.3 |
| DAG 22:6_22:6 | 5943 | 670251 | 0.99851 | 0.1-10000.0 | 0.279 | 0.1 | 0.3 |

### 3.3 Analysis of pure oil

The aim of this study was to determine whether DAG profiles of marine and plant oils can be used for the detection of adulteration with a performance comparable to TAG or PL-based analyses. Due to the simplicity of DAG profiling and the relatively short analysis time, this approach may provide a more accessible and cost-effective alternative for oil authentication.

To evaluate the differences in DAG profiles among oils of different origins, quantitative DAG analysis was performed on pure oil standards, including FO, KO, SFO, and SBO (*Suppl. Materials 1, Table 1)*.

PCA score plots (Figure 2iv.) demonstrated clear separation among all four oil groups, indicating distinct DAG composition profiles. The first principal component (PC1) explained 98.2% of the total variance and primarily separated plant-derived oils from marine-derived oils. The second principal component (PC2), accounting for 1.1% of the variance, contributed to further discrimination among the different oil groups. The observed separation reflects differences in fatty acid composition between marine and plant oils, particularly the presence of long-chain polyunsaturated fatty acids, such as EPA and DHA, in marine oils, which are absent or present at very low levels in plant oils. Replicates within each group clustered closely together, demonstrating strong intra-group consistency and high analytical reproducibility.

The fatty acyl composition of the DAG fraction was calculated from the quantitative DAG data. The molar content of each fatty acyl species was determined by summing the concentrations of all DAG molecular species containing the corresponding fatty acyl chain, multiplied by the number of occurrences of that fatty acyl chain within each DAG molecule by the following formula:

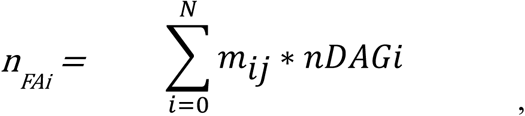

*where m – number of fatty acyls occurrences, n – is molar concentration*.

The molar percentage of each fatty acyl species was subsequently calculated to determine the relative fatty acyl composition of the oils. Figure 2 ii demonstrated the fatty acyls composition of DAG fraction from KO, FO, SFO and SBO. In KO, the most abundant fatty acyl species in DAG were 16:0 (22.3 ± 0.6%), 18:1 (15.2 ± 0.8%), 20:5 (23.6 ± 0.7%), and 22:6 (17.2 ± 1.2%).

The fatty acyl composition of the DAG fraction showed characteristics intermediate between those reported for the PL and TAG fractions by Xie et al., 2019. The PL fraction was dominated by 20:5, which accounted for approximately 30% of the total fatty acyl content, followed by 16:0 and 22:6 at approximately 23% and 15%, respectively. In contrast, the TAG fraction contained relatively higher proportions of 18:1 and 16:1 fatty acids.

A similar fatty acyl profile was observed for FO, although FO differed from KO by exhibiting a lower proportion of 18:1 and higher proportions of 14:0 and 16:1 fatty acyls. Overall, KO and FO exhibited highly similar fatty acyl compositions in their DAG fractions, suggesting that adulteration of KO with FO may be more challenging to detect compared to plant oils.

The plant-derived oils, SFO and SBO, predominantly consists of 18:1, 16:0, 18:2, and 18:3 fatty acyl species, but differ in their relative abundances of these fatty acyls. SFO contained a higher proportion of 18:1, whereas SBO was characterised by higher proportions of 16:0 and 18:3. These differences in fatty acyl composition contributed to the distinct DAG profiles observed between the two plant-derived oils.

The DAG profiles also exhibited distinct signatures among the pure oils analysed. Figure 2iii. shows a heatmap of the DAG species commonly detected across all oils, based on their absolute concentrations. In contrast, Figure 2i. shows the DAG composition of KO and FO, including all DAG species identified in the pooled marine oil samples. The heatmap in Figure 2 reveals that the oils are distinguished by differences in the relative abundances of individual DAG molecular species.

### 3.4 Adulteration of krill oil experiments

As shown by the fatty acyl profiles and DAG heatmaps, FO and KO exhibit highly similar DAG compositions. Therefore, the detection of FO adulteration in KO is expected to be challenging, particularly at low levels of adulteration.

KO samples adulterated with 0, 4, 10, and 20% FO were analysed (*Suppl. Materials 1, Table 2*) in terms of their DAG profiles using the developed method, and the resulting data were subjected to PCA (Figure 4i.). The pure KO samples (0% FO) were clearly separated from the adulterated samples and formed a distinct cluster. The adulterated samples were also grouped according to the level of FO addition, with the 4, 10, and 20% FO-adulterated samples showing progressive separation from one another. This clustering pattern indicates that the DAG profile was sensitive to the level of FO adulteration and that the developed method could discriminate KO samples according to the extent of adulteration. The increasing separation of the samples with increasing FO content suggests that the compositional changes in the DAG profile become more pronounced as the proportion of FO increases.

**Figure 4.**
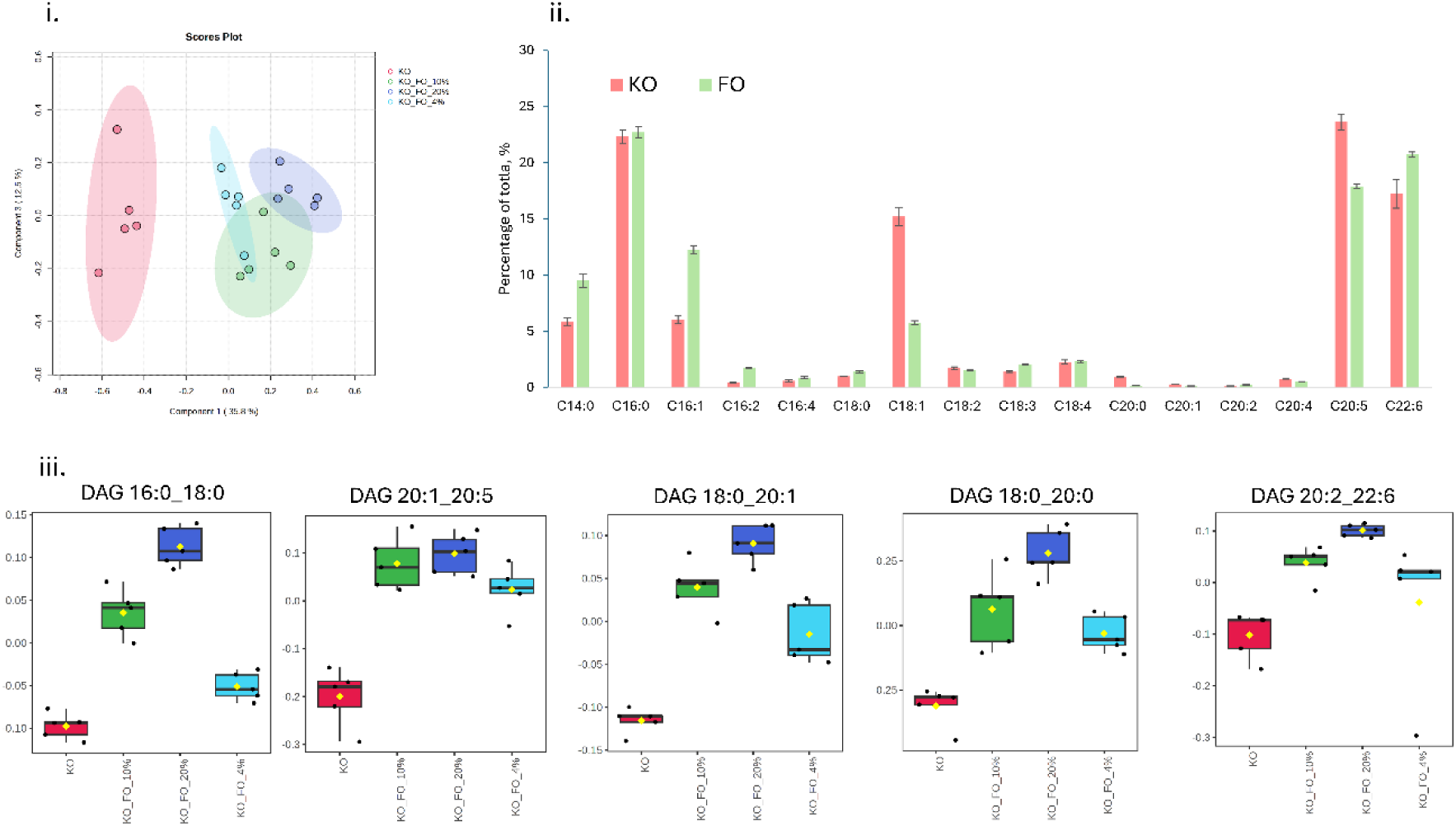
Detection of FO adulteration in KO based on DAG molecular profiles: (i) PCA score plot showing the separation of pure KO and KO samples adulterated with different percentages of FO. (ii) Fatty acyl composition of the DAG fraction in KO and FO samples calculated from DAG molecular species data. (iii) Box plots showing the distribution of DAG species selected by both PLS-DA and random forest analyses. Statistical significance was evaluated using one-way ANOVA followed by Tukey’s multiple comparison test.

To identify molecular features capable of discriminating FO-adulterated KO samples, the dataset containing was analysed using two complementary supervised approaches: partial least squares discriminant analysis (PLS-DA) and random forest classification (Schwarz et al., 2025). Random forest set up were as follows: number of trees to grow – 500, number of predictors to try – 7, randomness – on. Random forest classification achieved an out-of-bag error rate of 5%, corresponding to an overall classification accuracy of 95%. All KO, KO+10% FO, and KO+20% FO samples were correctly classified, while one KO+4% FO sample was misclassified as KO+10% FO. These results demonstrate the discriminative ability of the selected lipid features for detecting FO adulteration in KO samples.

Variables contributing most strongly to sample discrimination were ranked based on their variable importance in projection (VIP) scores from the PLS-DA model and variable importance values from the random forest model (*Suppl. Materials 2, Tables 1-2*). The 15 highest-ranked features from each model were compared, and only molecules consistently selected by both approaches were considered robust candidate markers of FO adulteration. This combined strategy was applied to minimize the influence of model-specific feature selection and to identify molecular markers with the greatest potential for reliable discrimination of adulterated KO samples.

The discriminatory potential of the selected markers was further evaluated by one-way ANOVA followed by false discovery rate (FDR) correction. The following DAG species were found in both lists: DAG 16:0_18:0, DAG 20:1_20:5, DAG 18:0_20:1, DAG 18:0_20:0, DAG 20:2_22:6 (Figure 4iii.). All selected markers showed significant differences among the sample groups after multiple-testing correction. Tukey’s post-hoc analysis was subsequently performed to identify the specific group differences responsible for the observed variation. As shown in (Figure 4iii.), the abundance of the selected markers directly correlated with increasing FO adulteration levels, with significant differences observed between KO and FO-adulterated samples. These results confirm the ability of the identified markers to discriminate KO samples adulterated with FO.

A separate experiment was conducted in which KO samples were adulterated with different levels (2.5, 5.0, 10.0, and 15.0%) of two plant oils, SFO and SBO. Figure 5ii. presents the PCA score plots obtained from the analysis of pure and adulterated KO samples, separately for SFO and SBO adulteration. In both cases, the pure KO samples were clearly separated from the adulterated samples, demonstrating that the DAG profile was sensitive to the presence of the added plant oils. Furthermore, a concentration-dependent separation pattern was observed, with the samples progressively shifting away from the pure KO cluster as the level of adulteration increased. The distinct clustering of samples according to the adulteration level indicates that the changes in the DAG profile were related to the proportion of added SFO or SBO. These results demonstrate that the developed method has the potential not only to discriminate pure KO from adulterated samples but also to differentiate samples according to the extent of adulteration.

When the pure KO and KO samples adulterated with either SBO or SFO were analysed together, the pure KO samples formed a distinct and closely grouped cluster, clearly separated from the adulterated samples (Figure 5i.). To further investigate the discrimination of adulterated samples and identify potential markers of adulteration, the data were subjected to random forest and PLS-DA analyses (*Suppl. Materials 2, Tables 3-4*), following a similar approach to that used for the FO adulteration study. The variables identified by both approaches are presented in Figure 5iii. The identified markers included DAG species originating from the added plant oils, as well as endogenous KO DAG species whose relative abundance decreased with increasing levels of adulteration. One example is DAG 20:2_22:6, which showed a progressive decrease with increasing adulterant concentration. The complementary changes in DAG species, including the increase in plant-oil-derived DAGs and the decrease in characteristic KO DAGs, provided a clear molecular signature of adulteration and contributed to the discrimination of pure and adulterated KO samples.

**Figure 5.**
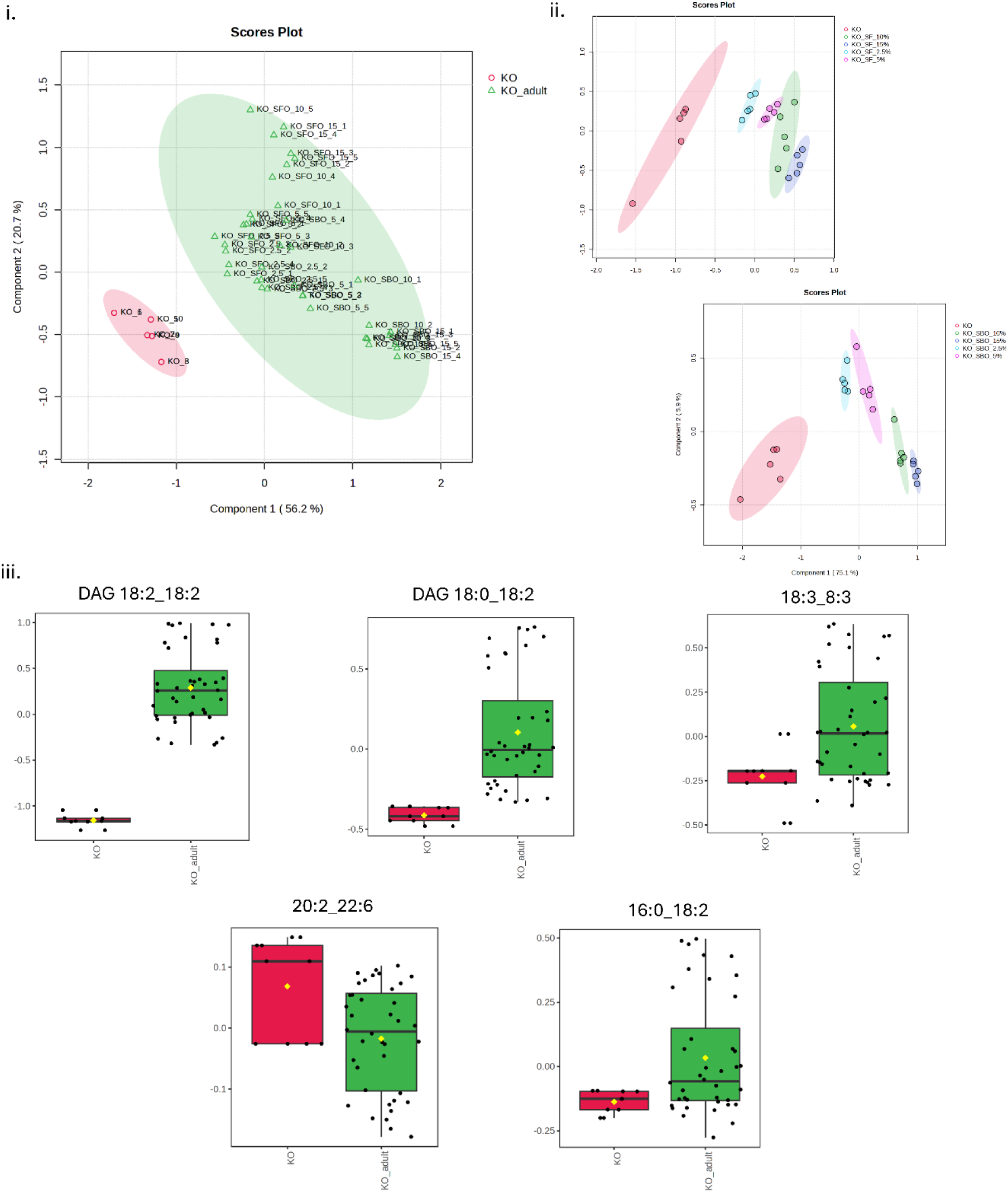
Detection of FO adulteration in KO based on DAG molecular profiles: (i) PCA score plot showing the separation of pure KO and KO samples adulterated with different percentages of SBO and SFO. (ii) PCA score plot showing a separate analysis of KO samples adulterated with SBO and SFO. (iii) Box plots showing the distribution of DAG species selected by both PLS-DA and random forest analyses. Statistical significance was evaluated using one-way ANOVA followed by Tukey’s multiple comparison test.

## Conclusion

A rapid and comprehensive HPLC-MS/MS method based on multiple reaction monitoring (MRM) was successfully developed for the analysis of DAG molecular species in plant and marine oils. The method enabled the determination of 75 DAG molecular species within 5 min, with a six-fold linear range and low limits of detection and quantification, providing an efficient approach for DAG profiling. Its applicability was demonstrated through the detection and discrimination of KO adulterated with FO, SBO, and SFO. PCA showed clear separation between pure and adulterated KO samples, as well as concentration-dependent changes associated with increasing adulterant levels. Furthermore, the combined use of PLS-DA and random forest analysis enabled the identification of potential DAG markers characteristic of FO, SBO, and SFO adulteration. Overall, the developed method provides a rapid, sensitive, and informative analytical approach for DAG profiling and supports the detection and characterisation of adulteration in marine and plant oils.

## CRediT authorship contribution statement

Daria Makeeva: Methodology, Validation, Visualization, Investigation, Formal Analysis, Data Curation, Software, Writing – Original Draft. Louise V. Michaelson: Conceptualization, Investigation, Methodology, Writing – Review and Editing, Resources, and Funding Acquisition.

## Acknowledgment

The authors acknowledge the use of instruments at the Lipidomic and Metabolomics facility of Rothamsted Research, United Kingdom.

## Supplementary data

Supplementary materials 1: Tables 1-4 (Supplementary material_1.xlsx). Supplementary material 2: Tables 1-4, Figure S1 (Supplementary material_2.xlsx).

## Funding Sources

This work and authors were supported by the BBSRC (UK) funded Institute Programme Grant Green Engineering, BB/X010988/1.

## Notes

### Competing Interest Statement

The authors have declared no competing interest.

